# Senescence gives insights into the morphogenetic evolution of anamniotes

**DOI:** 10.1101/091199

**Authors:** Éric Villiard, Jean-François Denis, Faranak Sadat Hashemi, Sebastian Igelmann, Gerardo Ferbeyre, Stéphane Roy

## Abstract

We report the presence of senescent cells in several transient structures in developing amphibian and teleost fish, suggesting novel mechanisms of morphogenesis that appeared early in vertebrate evolution.

**Abstract:** Senescence represents a mechanism to avoid undesired cell proliferation that plays a role in tumor suppression, wound healing and embryonic development. In order to gain insight on the evolution of senescence, we looked at its presence in developing axolotls (urodele amphibians) and in zebrafish (teleost fish), which are both anamniotes. Our data indicate that cellular senescence is present in various developing structures in axolotls (pronephros, olfactory nerve fascicles, lateral organs, gums) and in zebrafish (epithelium of the yolk sac and in the lower part of the gut). Senescence was particularly associated with transient structures (pronephros in axolotls & yolk sac in zebrafish) suggesting that it plays a role in the elimination of these tissues. Our data supports the notion that cellular senescence evolved early in vertebrate evolution to influence embryonic development.

## Manuscript

Cellular senescence has been, until recently, linked to aging and as a mean to prevent aberrant cellular proliferation. Cells that become senescent stop proliferating and secrete multiple cytokines that can act as growth factors and immunomodulators. Although senescence per se is not cell death, senescent cells *in vivo* are cleared by phagocytosis (Xue et al., 2007). The initial view of senescence as a mean to eliminate premalignant cells has been radically changed by the groups of Serrano and Keyes who independently published that cellular senescence is also important for embryonic development in mice (Munoz-Espin et al., 2013; Storer et al., 2013). More recently, Yun et al published that senescent cells were transiently present in regenerating limbs of axolotls where they are eliminated by macrophages (Yun et al., 2015). These observations provide new insight on the importance of cellular senescence during development. To determine how widespread developmental senescence is, we examined senescence in axolotl and zebrafish embryonic development.

Developing axolotl and zebrafish embryos were fixed glutaraldehyde and stained for the Senescence Associated β-Galactosidase (SaβG). We detected a strong SaβG activity in the pronephros of axolotls (fig.1 A, B, H, I). SaβG activity was also detected along the pronephric ducts from the pronephros all along until the bladder (fig.1 B). Some primitive fishes retain pronephros all their life but in salamanders it is present transiently and is replaced by the mesonephros as they reach maturity (Hildebrand, 1988). A second strong staining area for senescent cells was detected in the olfactory nerve fascicles in the nostril area (fig.1 A & E). If the SaβG assay was allowed to proceed for an extra 2-4 hours at 37°, other structures appeared as possibly containing senescent cells. This is the case of the roof plate of the developing spinal cord (fig. 1 C, H) and the gum cells just adjacent to protruding teeth (fig. 1 D & E). Interestingly, the latter observation of senescent cells at the point where teeth are erupting through the gums are reminiscent of the mammalian enamel knot. Cells comprising the enamel knot do not proliferate and end up being eliminated by apoptosis to allow the teeth to grow through the gum epithelium (Matalova et al., 2004). The enamel knot is a signalling center, similar to the apical ectodermal ridge (aer) of the amniote limb bud. Storer et al demonstrated that aer cells are positive for senescence even though they also display apoptosis (Francis et al., 2005; Storer et al., 2013). Finally, we observed SaβG activity associated with melanocytes in the skin (fig. 1 F) and in the lateral organs (fig.1 C, G & H).

**Fig.1:**
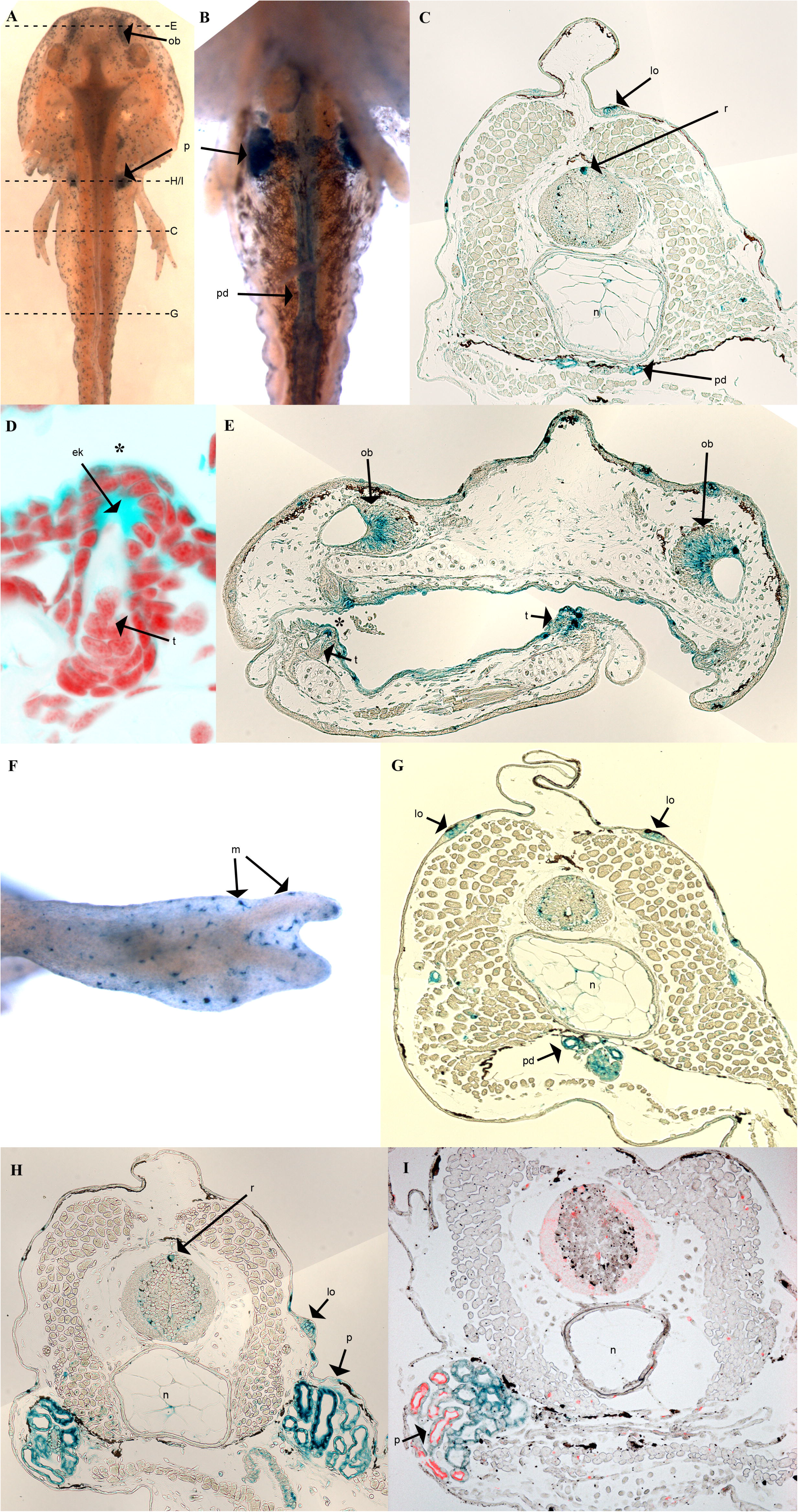
Axolotl stage 50, X-gal stained for senescence (SaβG): (A) Dorsal view, showing subsequent transversal sections. (B) Ventral view. (C) Mid body transverse section. (D) Teeth, magnified from panel E, marked with *. (E) Rostral transverse section. (F) Fore limb. (G) Caudal transverse section. (H) Transverse section of pronephric area. (I) Transverse section of pronephric area double stained for SaβG (blue) and phospho-*Erk1/2* (red). *onf*: olfactory nerve fascicle; *p*: pronephros; *pd*: pronephric duct; *lo*: lateral organ; *r*: roof plate; *n*: notochord; *ek*: enamel knot; *t*: teeth; *m*: melanocytes. Number of replicates (each n represents an animal) for A and B n=10; for panels C – H n=5; and panel I n=3.

Senescence in mammals is often associated with the presence of phospho-Erk (Deschenes-Simard et al., 2013). We used the phospho-Erk1/2 antibody to assess whether it was present in the axolotl pronephros (strongest staining organ). Our results show that phospho-Erk1/2 is indeed present in the axolotl developing pronephros. This indicates that signals associated with cellular senescence are highly conserved across a vast evolutionary distance among vertebrates from amphibians to mammals.

In order to further assess the evolutionary importance of senescence developing zebrafish were stained for SaβG activity. SaβG staining is detected at very early stages including the yolk sac (fig. 2), a temporary structure used as a source of energy for the developing embryo that is lost over time (Kimmel et al., 1995). Our results suggest that senescence of this structure is partly responsible for its elimination. The earliest detection of senescence in zebrafish is at the 32-64 cell stage (fig. 2L). At the 8-13 somite stage senescence was detected in the yolk sac (fig. 2J). From the 20-25 somite stage until 4 days post-fertilization senescence is detected in the yolk sac and in the tissues that will become the intestine (fig. 2E-I). From day 7 until day 15 senescence became restricted the cloacal end of the intestine (fig. 2A-D). We did not detect any senescence signal in the zebrafish’s pronephric area which may be due to the fact that bony fish have rudimentary kidneys all their life and therefore there is no need to eliminate them. Eliminating structures using cellular senescence provides opportunity for the surrounding tissues to replace or compensate the loss of those cells, which would benefit the morphogenetic process. In addition, the products secreted by senescent cells can stimulate cellular growth and migration of adjoining cells and thus have an impact on development.

**Fig.2:**
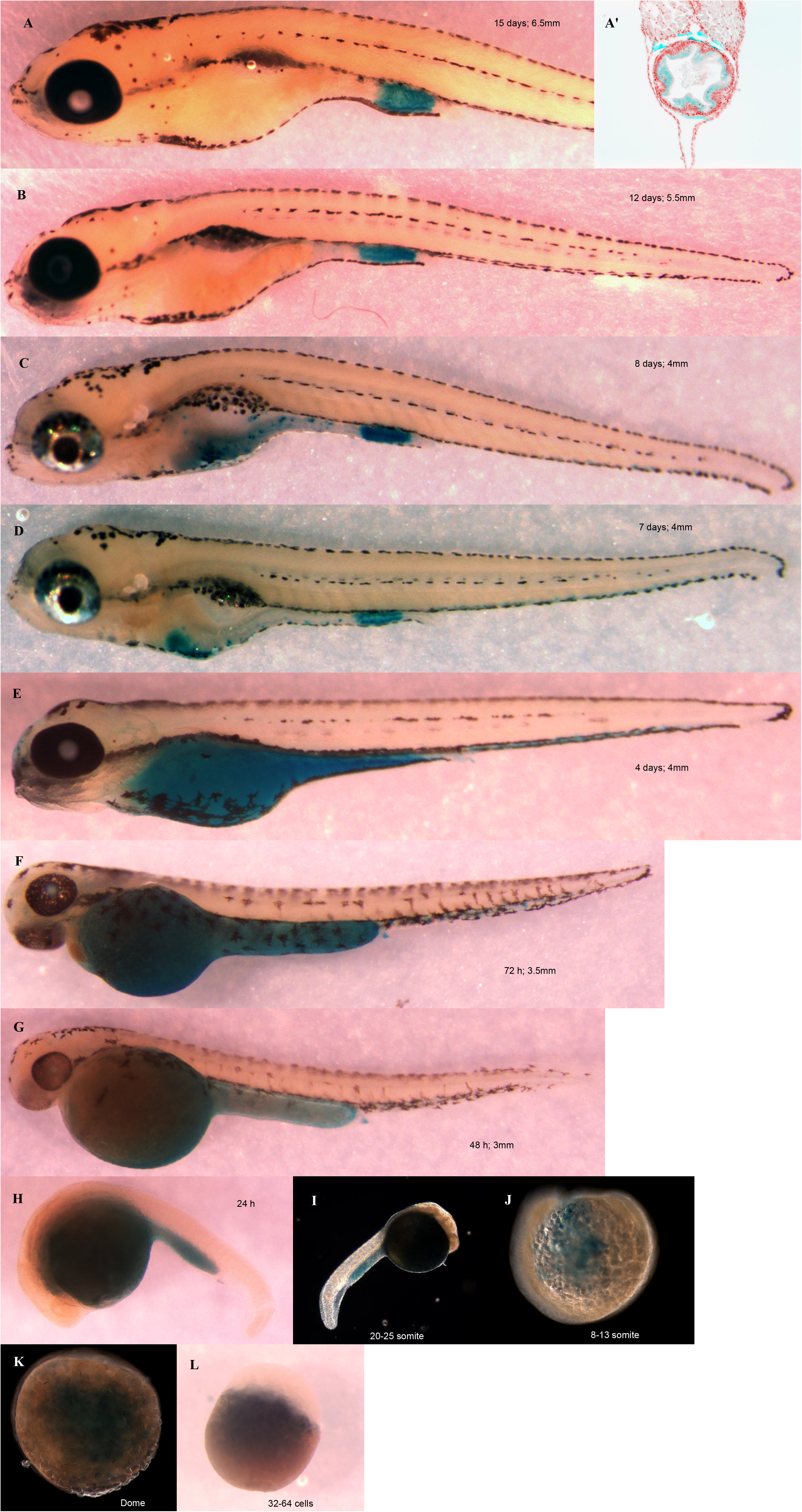
Zebrafish stages, X-gal stained for senescence (SaβG): Presented from (A) 15 days post-fertilization animal; (A’) transverse section of a 15 days post-fertilized animal, cloaca level, Dapi staining of nuclei (red) and SaβG staining (blue); (B) 12 days; (C) 8 days; (D) 7 days; (E) 4 days; (F) 72 hours; (G) 48 hours; (H) 24 hours; (I) 20-25 somite; (J) 8-13 somite; (K) Dome; and (L) 32-64 cells embryo. The length of the embryos is indicated in mm on each panel. Number of replicates (each n represents an embryo) is n=8.

The discovery of senescence during embryonic development does not support the idea that it evolved for the sole purpose of protecting organisms from oncogenic transformation. It is more likely that it evolved primarily for embryonic development and is re-utilized in mature animals as a mechanism of tumor suppression. This report demonstrates that cellular senescence is present during the embryonic development of lower vertebrates (salamander and teleost fish). The paper by Yun’s group (this issue) also shows that senescence is present in Xenopus as well as in axolotls. Our results combined with the work by the groups of Yun, Serrano and Keyes (Munoz-Espin et al., 2013; Storer et al., 2013) clearly demonstrate that senescence is a cellular process that evolved early in the vertebrate genealogy as it is present in organisms ranging from bony fish to amphibians all the way to mammals. The novel observation that cellular senescence is a bona fide process of embryogenesis opens the door for new investigations on the evolutionary pressures shaping development and raises important questions about the evolution of molecular pathways for the selective induction of senescence during embryonic development. A developmental control of cellular senescence raises the intriguing possibility that the observed increase of senescent cells during aging could be developmentally programmed as well.

## Materials and Methods

### Senescence Associated β-Galactosidase

Embryos were fixed with 0.5% glutaraldehyde in PBS at 4☐ C 6 hours to overnight and then rinsed 2 times 15 min in PBS pH5.5 containing 1mM MgCl_2_ at 4☐ C. The embryos were stained for SaβG in X-gal staining solution (0.1% X-gal, 5 mM potassium ferrocyanide, 5 mM potassium ferricyanide, 150 mM Sodium chloride, and 2 mM Magnesium chloride in PBS, pH 5.5) 4-6 hours at 37☐ C. Controls to confirm SaβG staining specificity were performed in identical conditions except the pH of the solutions were adjusted to 7.5 for which SaβG is inactive (none of the controls showed any sign of β-gal activity, n=10).

### Immunofluorescence enhanced with Tyramide

Sections were rehydrated as previously described then blocked using 2% BSA in TBS-T for 1h at room temperature (Levesque et al., 2010). Primary antibody was diluted in blocking solution (anti-p-Erk1/2, cat# 4370, cell signalling, 1/400) and incubated on slide overnight at 4°C. Secondary HRP coupled antibody was diluted (anti-rabbit HRP, cat# 170-6515, Biorad, 1/800) in blocking solution for p-Erk1/2 and incubated at room temperature for 45 min. Tyramide (Biotium, San Francisco Bay, CA, cat# 92175) was diluted in 1X TBS with 0.0015% H_2_O_2_ to an active concentration of 11.6 µM then incubated at room temperature for 8 min. All slides were mounted with ProLong® Gold antifade reagent containing DAPI (Invitrogen, cat# 36931). Slides were visualized with a Zeiss Axio Imager M2 Optical Microscope (Zeiss, Munich, Germany).

### Animal ethics conformity

Axolotl wild type embryos were obtained from mating adult axolotls at the Université de Montréal. Zebrafish were a generously provided by the laboratory of Dr Moldovan from the Sainte-Justine Hospital Research Institute affiliated with the Université de Montréal. All experiments and manipulation of animals were done in accordance with the requirement of the Université de Montréal animal ethics committee which is overseen by the Canadian Council of Animal Care.

## Acknowledgements

We acknowledge the help of Drs Moldovan and Lévesque for providing wild type zebrafish. JF Denis is supported by scholarships from the Réseau de Recherche en Santé Buccodentaire et Osseuse and the Faculty of Graduate Studies. F Hashemi was the recipient of a summer studentship from the Fond Ernest Charron, Faculty of Dentistry. GF is a chercheur national FRQS.

## Author contributions

EV performed over 60% of the experiments and helped analyze/interpret the data and he co-wrote the manuscript. JFD helped with 25% of the experiments. FSH and SI helped with some of the animal treatments. GF and SR designed the project and supervised it and they helped analyze/interpret the data and co-wrote the manuscript.

